# Actigraphy-Based Movement Profiles and Their Association With Circadian Rhythms Integrity in Real-World Settings

**DOI:** 10.64898/2025.12.19.695124

**Authors:** Mariana Marchesano, Ana Silva, Bettina Tassino

## Abstract

Both active movement profiles and robust circadian rhythms are linked to improved health outcomes, yet the underlying mechanisms remain partially understood. We investigated this relationship in young adults (n = 169, aged 18–30 years) under real-world conditions using actigraphy data. We performed k-means clustering on 12 accelerometer-based features capturing magnitude, duration, frequency, and intensity distribution to derive movement behavior profiles. As a proxy of circadian rhythms integrity we computed the Circadian Function Index (CFI), which combines intradaily variability, interdaily stability, and relative amplitude. We also assessed circadian phase and sleep quality parameters. Additionally, we quantified light exposure and physical activity over 3-hour daily intervals. The unsupervised algorithm identified two non-overlapping profiles among participants, the More Active (MA) and the Less Active (LA) profiles. MA exhibited a higher CFI (0.81 ± 0.06 vs. 0.69 ± 0.06, *p* <0.001), which was also positively associated with early-evening physical activity, but not with light exposure. MA also showed an earlier activity-based phase indicator, the midpoint of the five least active hours (L5c, 04:30 ± 01:03 vs. 04:59 ± 01:15, *p* adj. = 0.04), which was inversely associated with early-morning physical activity and late-morning light exposure. We found no differences in sleep quality between MA and LA. Our results underscore the association between movement behavior and overall circadian rhythms integrity. Importantly, these findings reinforce actigraphy as a multidimensional tool for both health research and clinical applications.

---

The health benefits of physical activity are well established, with global guidelines recommending specific weekly frequencies, intensities, and durations according to age (Bull et al., 2020), although the precise underlying mechanisms remain debated (Chow et al., 2022; Neufer et al., 2015). Growing evidence highlight the relevance of timing in modulating these effects (Bennett and Sato, 2023); however, current guidelines still offer no indications on *when* to exercise (Kim et al., 2023). Circadian rhythms—endogenous oscillations with a period of approximately 24 h—are entrained to environmental cycles, enabling anticipation of periodic events and tuning physiological, behavioral, and metabolic processes (Paranjpe and Sharma, 2005). Since movement behavior is essential for development and survival, it is expected to be tightly linked to biological timing mechanisms. Although hard to disentangle, this relationship unfolds in two directions: circadian rhythms influence physical activity performance, for example, by accounting for the diurnal variation in glucose and lipid metabolism, while physical activity contributes to the regulation of the circadian system, particularly through its effects on the skeletal muscle clock (Gabriel and Zierath, 2019; Kim et al., 2023). In both directions, this interplay may affect health and life quality, and has consequently become an increasingly active area of research (Drăgoi et al., 2024).

Exercise has only recently been recognized as a genuine entrainer of the human circadian system (Lewis et al., 2018; Youngstedt et al., 2019). Specifically, strong evidence shows that early-morning and afternoon scheduled exercise induce phase advances, whereas nighttime exercise elicits phase delays, independently of light exposure —the primary *zeitgeber*, or time-giver (Coirolo et al., 2022). In urban context, where the photic signal has been weakened, with consequent adverse impacts on health outcomes (Roenneberg and Merrow, 2016), exercise emerges as a promising non-pharmacological strategy for the prevention and treatment of circadian disorders (Shen et al., 2023).

Healthy circadian rhythms are expected to be regular, non-fragmented, robust and in phase with the light-dark cycle (Van Someren and Riemersma-Van Der Lek, 2007). At the molecular level, exercise re-sets clock genes in skeletal muscle and other tissues, affecting both the amplitude and phase of circadian rhythms (Gabriel and Zierath, 2019; Zambon et al., 2003). Based on both clinical and preclinical evidence, exercising at specific times of day has been proposed as a therapeutic strategy to improve cardiometabolic health (Martin et al., 2023; Procopio and Esser, 2025), and has been associated with a reduced risk of cancer (Weitzer et al., 2021), and lower all-cause mortality (Bennett and Sato, 2023). These benefits may also be influenced by the intensity, duration, frequency, and type of exercise (Shen et al., 2023).

Wearable devices provide a non-invasive way to objectively assess circadian and physical activity patterns, increasingly proving valuable in clinical and research settings (Patterson et al., 2023). Classical parametric approaches involve fitting data to a cosinor model (Gubin et al., 2025; Halberg et al., 1967); however, under real-life conditions, non-parametric proxies of circadian phase—such as the midpoint of the five least active hours (L5c)—often provide more reliable estimates in the presence of irregular activity patterns and data noise (Gubin et al., 2025; Van Someren et al., 1999). Other three key circadian parameters are interdaily stability (IS), intradaily variability (IV), and relative amplitude (RA), which respectively account for day-to-day regularity, within-day fragmentation, and the consolidation of daytime activity and nighttime rest (Gubin et al., 2025; Witting et al., 1990). Interestingly, they can be integrated into a unique value to quantify the overall integrity of the circadian activity–rest rhythm through the Circadian Function Index (CFI, Ortiz-Tudela et al., 2010). Ranging from 0 (absence of circadian rhythmicity) to 1 (highly robust circadian rhythm), the CFI has been applied in clinical and comparative studies, with healthy controls consistently exhibiting higher values (Madrid-Navarro et al., 2018; Martinez-Nicolas et al., 2021). Regarding physical activity, Yerramalla et al. (2024) recently identified daily movement behavior profiles in older adults using accelerometer-derived features and an unsupervised approach, demonstrating that the most active profile was associated with the lowest all-cause mortality risk. However, the relationship between movement profiles and circadian rhythms integrity, as indexed by the CFI, remains unexplored.

In recent years, translational chronobiology has increasingly sought non-invasive behavioral indicators of circadian health as well as the use of physical activity as therapeutic strategy for circadian re-entrainment (Klerman et al., 2022; Shen et al., 2023). In this study, based only on actigraphy data, we test the hypothesis that persons with more active movement profiles will exhibit higher circadian rhythms integrity than those with less active profiles. We examine the association of movement profiles—derived using an unsupervised algorithm —with actigraphic indicators of circadian rhythms integrity, circadian phase, and sleep quality in healthy young adults under real-world conditions.

## METHODS

### Participants

A total of 95 Uruguayan young adults (18-30 years old), without prior psychiatric, neurological, or sleep diagnoses, or taking sleep-affecting medications, were recruited between September and October 2024, and provided written consent. All study procedures were approved by the Ethics Committee of the School of Psychology, Universidad de la República (№ 191175-000124-23), and complied with the principles of the Declaration of Helsinki (World Medical Association, 2013). In addition to these newly enrolled participants, the dataset incorporated records from previous studies conducted under comparable protocols. These studies were carried out during peri-equinox periods, in March 2016 (*n* = 13, Castillo et al., 2023; Silva et al., 2019), September 2019 (*n* = 30, Coirolo et al., 2020, 2022), and September 2021 (*n* = 31, Marchesano et al., 2023, 2025). The final sample included 166 participants (Table 1). In all cases, chronotype was assessed as the midsleep point on free days corrected for sleep debt on workdays (MSFsc, Roenneberg et al., 2004) using the Spanish version of the Munich Chronotype Questionnaire (MCTQ, Roenneberg et al., 2003).

**Table 1.** Sociodemographic and chronobiological data.

| Data Sample | Sex (Female) | Age (Years) | MSFsc (h) | N |
| --- | --- | --- | --- | --- |
| University students (new data) | 65% (62) | 23.4 (3.6) | 05:13 (01:55) | 95 |
| Dancers (Marchesano et al., 2025) | 87% (27) | 22.5 (3.4) | 05:13 (01:22) | 31 |
| Dancers (Coirolo et al., 2022) | 80% (24) | 22.4 (2.8) | 06:07 (01:51) | 30 |
| University students (Silva et al., 2019) | 69% (9) | 22.8 (1.5) | 05:57 (00:50) | 13 |
| **Overall** | 72% (122) | 23.0 (3.3) | 05:25 (01:48) | 169 |

Data are given as % (*n*) or mean (SD).

### Actimetry

All participants—including those newly recruited and those from previous studies—were equipped with GeneActive Original (Activinsights) tri-axial accelerometers to wear on their non-dominant wrist for a minimum of 7–17 days, depending on the cohort, programmed to record at 10 Hz. This instrument allowed us to obtain sleep and activity patterns, physical activity levels, and light exposure. Activity was estimated using the Euclidean Norm Minus One (ENMO) (Van Hees et al., 2013) and expressed in gravity-based acceleration units (g), where 1 g = 9.81 m·s^−2^ (Hildebrand et al., 2014). Values were then converted to milligravity units (mg; 1 g = 1000 mg). We processed data with the GGIR package (Migueles et al., 2019) in the RStudio environment (Posit team, 2025; R Core Team, 2025). We characterized circadian and 24 h variability of activity using both parametric and non-parametric metrics (Gubin et al., 2025). Parametric rhythmicity was assessed through Cosinor analysis, from which we estimated amplitude, mesor, and acrophase (Cornelissen, 2014; Halberg et al., 1967). Non-parametric analysis (Van Someren et al., 1999; Witting et al., 1990) provided complementary parameters: M10, the average of the 10 hours with the highest values; M10c, the midpoint of the M10 period; L5, the average of the 5 hours with the lowest values; L5c, the midpoint of the L5 period; RA, relative amplitude; IV, intra-daily variability; and IS, inter-daily stability.

RA (relative amplitude) reflects the difference between M10 activity and L5 activity, with values ranging from 0 to 1, where higher values indicate greater daytime activity and reduced activity during sleep (Van Someren et al., 1998). It was calculated as follows:

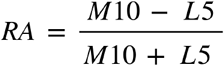

IV (intra-daily variability) quantifies the variability of activity within a day and can range from 0 to 2 (or exceed 2 if an ultradian component is present). Higher values indicate a more fragmented rhythm, reflecting shorter periods of rest and activity rather than a single extended active period during the daytime and a single extended rest period at night (Witting et al., 1990). IV was calculated by the following formula, where N—total number of measurements in the full time series, Xi—individual values at time i, Xm—mean of all Xi values (Gubin et al., 2025):

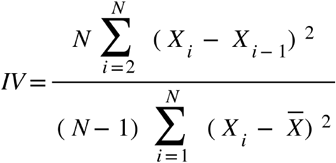

IS (inter-daily stability) measures how consistent the rest-activity pattern is between days and ranges from 0 to 1 (Witting et al., 1990). Values closer to 1 mean more constant rest-activity patterns. IS was calculated by the following formula, where N—total number of measurements in the overall time series, p—number of data points per 24 h, Xi—individual values at time i, X—mean of the overall time series, Xh—hourly means (Gubin et al., 2025):

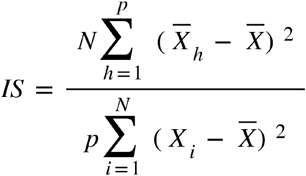

Circadian Function Index assesses the overall integrity of the circadian activity–rest rhythm, incorporating IV, IS and RA, with values ranging from 0 to 1, 1 being perfectly regular and robust rhythms, unfragmented, and with high amplitude, and 0 indicating absence of circadian rhythmicity (Ortiz-Tudela et al., 2010). IV values are inverted and normalized between 0 and 1, with 0 being a noise signal, and 1 a perfect sinusoid.

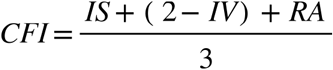

As validated elsewhere, acrophase, L5c, and M10c were considered as proxies of circadian phase (Bonmati-Carrion et al., 2014; Mitchell et al., 2017; Ruiz et al., 2020). To account for potential seasonal effects, calendar day length was calculated for each day of actigraphy recording as the difference in hours between sunrise and sunset, and then averaged across days for each participant. Sunrise and sunset times were obtained using the getSunlightTimes() function from the suncalc R package (Thieurmel and Elmarhraoui, 2017).

### Movement behavior features

Based on Yerramalla et al. (2024), we selected 12 accelerometer-derived features to capture 5 dimensions of movement behavior: overall activity level (mean per day, mg), total duration (min), frequency (number of bouts), bout duration (also reflecting fragmentation) (Diaz et al., 2017; Rowlands et al., 2018), and intensity distribution across the day (gradient and intercept) (Rowlands et al., 2018). The first four dimensions were assessed across three intensity levels—inactive (IN, PA ≤ 45.8 mg ), light (LPA, 45.8 < PA < 93.2 mg) and moderate-to-vigorous (MVPA, PA ≥ 93.2 mg) (Hildebrand et al., 2014).

### Sleep parameters

As indicators of sleep quality we used sleep duration, Sleep Efficiency, the Wake After Sleep Onset (WASO), and the Sleep Regularity Index (SRI). The recommended sleep duration for this age group ranges between 7 and 9 hours. Sleep efficiency (SE, %) was calculated for each night as the ratio of total sleep time to the main sleep period time, both derived from the HDCZA algorithm in GGIR (Van Hees et al., 2018), and multiplied by 100 (Sansom et al., 2023). WASO refers to the total time spent awake after the initial sleep onset (Cole et al., 1992). The SRI quantifies the regularity of sleep–wake patterns across consecutive days (Phillips et al., 2017) and ranges from −100 to 100, where 100 indicates perfect regularity (identical days), 0 indicates a random pattern, and −100 indicates perfectly reversed regularity.

### Clustering

We identified activity profiles following the procedure of Yerramalla et al. (2024), using standardized accelerometer-assessed features not related to timing (Figure 1). We applied the k-means clustering algorithm (via the kmeans() function in R), a method designed to maximize intra-class similarity while minimizing inter-class similarity, with the Hartigan-Wong algorithm (Hartigan and Wong, 1979). We determined the optimal number of clusters using the NbClust()function (Charrad et al., 2014), which evaluates multiple indices, including the elbow method, as documented by Kassambara and Mundt (2017), and the gap statistic (Tibshirani et al., 2001). We assessed clustering quality using silhouette analysis (Rousseeuw, 1987), computed with the silhouette() function from the cluster package in R (Maechler et al., 2025), which quantifies how similar an observation is to its own cluster relative to other clusters, with values ranging from _−_1 (poor clustering) to +1 (well-clustered).

**Figure 1.**
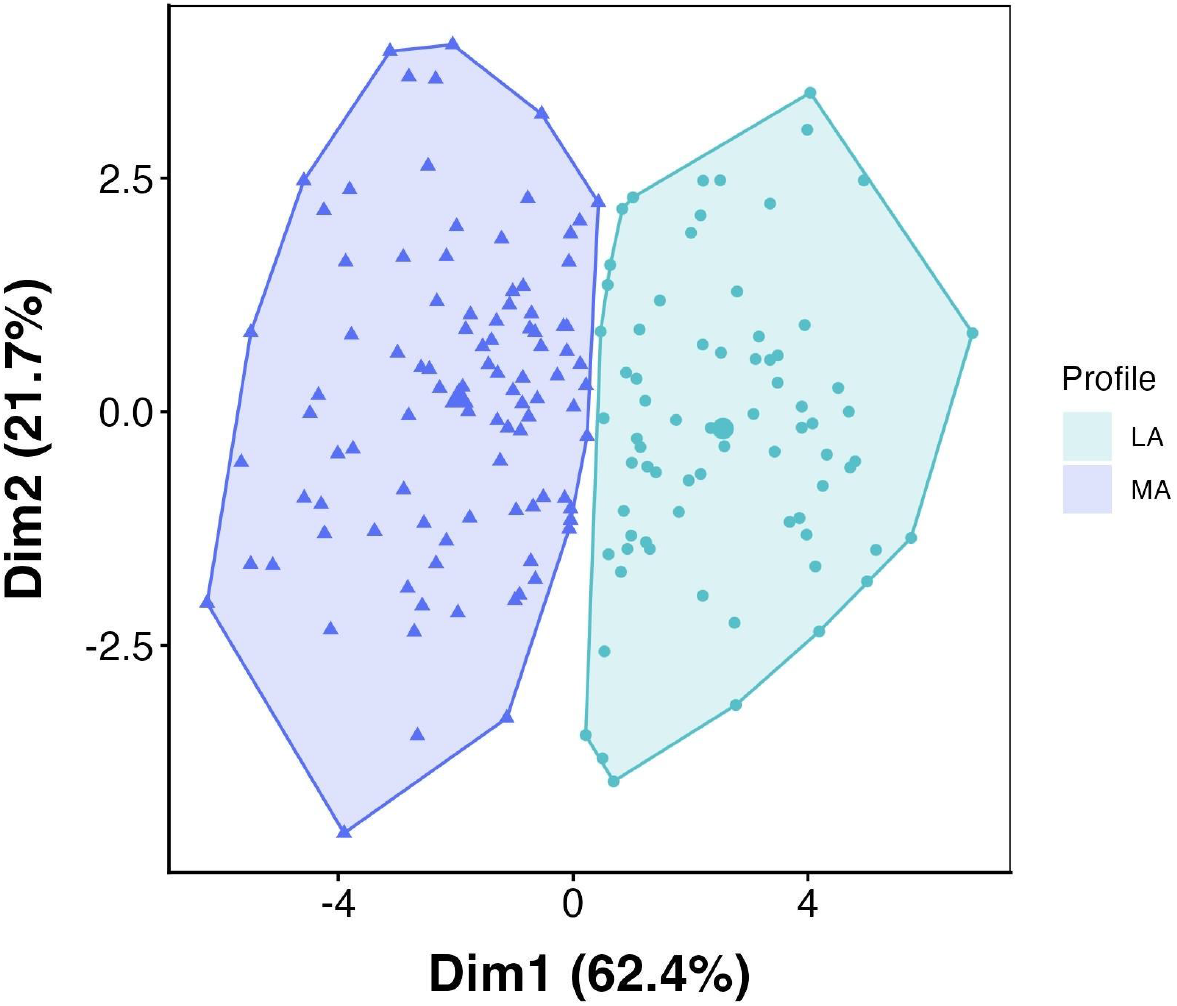
Movement behavior cluster analysis. K-means algorithm was used to identify profiles based on 12 features derived from accelerometry. The two cluster solution classified the data into a More Active profile (MA) and a Less Active profile (LA). MA was characterized by higher average daily acceleration, more daily minutes spent in light-intensity physical activity (LPA) and moderate-to-vigorous physical activity (MVPA), fewer minutes of inactivity, and more frequent and larger MVPA bouts. It also showed a lower intensity slope and intercept, indicating that individuals accumulated more time in higher-intensity activities compared with LA. Dim 1 and Dim 2 correspond to the projections onto the first two principal components (percentages indicate the variance explained). Each point represents one participant.

### Data analysis

We conducted all statistical analyses using R statistical software, version 4.5.1 (2025-06-13), within the RStudio environment (version 2025.9.1.401) (Posit team, 2025; R Core Team, 2025). Differences between profiles were assessed using Wilcoxon tests (Table 2, Figure 2, Table 3). Mixed-effects regression models, including individuals as random effects, were fitted to estimate the association of light exposure or accelerometry measures with time-of-day intervals, profile, and type of day (Figure 3). Marginal means were compared using the emmeans package (Lenth and Piaskowski, 2017) and multiple comparisons were adjusted using Holm–Bonferroni method (Holm, 1979). Statistical significance was set at *p*≤0.05. Linear regression models were fitted to examine the association between circadian parameters (CFI and L5c) and daily activity-light patterns. Predictor variables included the identified activity profiles and standardized, weighted measures of activity and light exposure for each time-of-day interval (Table 4). Lux values were log_10_-transformed prior to metric calculation using the transformation log_10_(Light + 1), in order to avoid undefined values caused by zeros (Wallace et al., 2025).

**Table 2.** Movement features by profile.

| Movement Features | LA<br>N = 73 | MA<br>N = 96 | *p*-Value adj | Effect Size |  |
| --- | --- | --- | --- | --- | --- |
| Average daily acceleration (mg) | 36 (6) | 56 (10) | <0.001 | 0.8 | Large |
| Total duration in IN (min/day) | 742 (53) | 615 (65) | <0.001 | 0.8 | Large |
| Total duration in LPA (min/day) | 147 (30) | 199 (31) | <0.001 | 0.7 | Large |
| Total duration in MVPA (min/day) | 89 (29) | 172 (45) | <0.001 | 0.8 | Large |
| Mean duration of IN bouts (min) | 17.9 (4.0) | 10.2 (2.1) | <0.001 | 0.8 | Large |
| Mean duration of LPA bouts (min) | 2.55 (0.31) | 2.48 (0.26) | 0.12 | 0.1 |  |
| Mean duration of MVPA bouts (min) | 3.27 (0.79) | 4.04 (0.84) | <0.001 | 0.4 | Moderate |
| Daily bouts in IN (*n*) | 45 (8) | 63 (8) | <0.001 | 0.7 | Large |
| Daily bouts in LPA (*n*) | 52 (10) | 75 (10) | <0.001 | 0.8 | Large |
| Daily bouts in MVPA (*n*) | 26 (6) | 43 (7) | <0.001 | 0.8 | Large |
| Intensity gradient | -1.96 (0.13) | -1.80 (0.10) | <0.001 | 0.6 | Large |
| Intensity intercept | 12.02 (0.44) | 11.65 (0.40) | <0.001 | 0.4 | Moderate |

Values are given as mean (SD). Wilcoxon rank-sum test. Bold-formatted *p*-values indicate statistical significance after Holm-Bonferroni.
correction. Abbreviations: IN=inactivity; LPA=light-intensity physical activity; MVPA=moderate-to-vigorous physical activity.

**Table 3.**
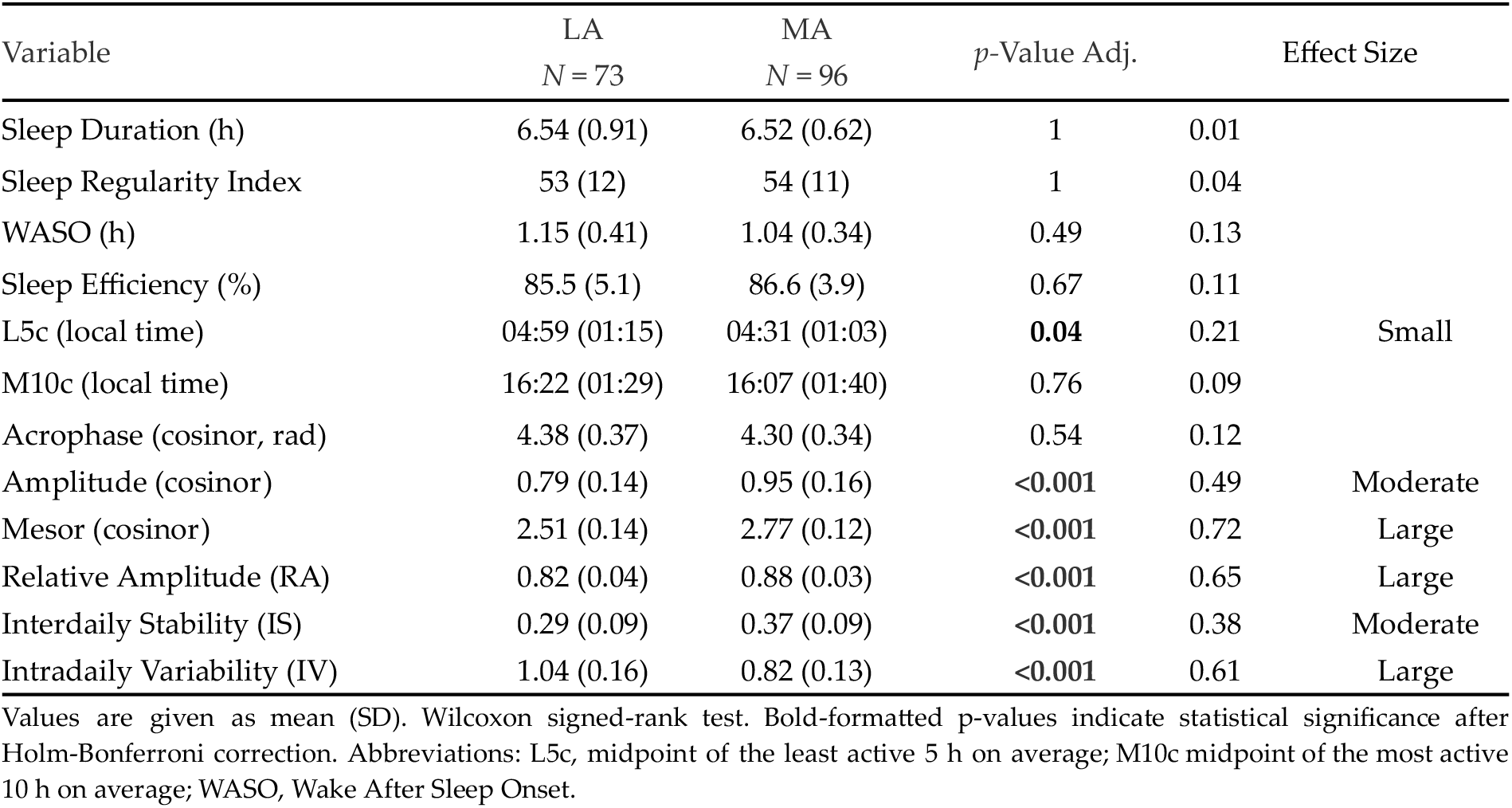
Circadian phase, circadian activity rhythms and sleep quality by movement profile.

**Table 4.** Associations between L5c CFI, profile and time of day for light exposure and activity.

| Predictors | L5c |  |  |  | CFI |  |  |  |
| --- | --- | --- | --- | --- | --- | --- | --- | --- |
| | B (SE) | $\beta$ | CI | $p$ | B (SE) | $\beta$ | CI | $p$ |
| (Intercept) | 4.66 (0.08) | -0.05 | 4.50 to 4.83 | **<0.001** | 0.79 (0.01) | 0.35 | 0.77 to 0.80 | **<0.001** |
| profile [Less Active] | 0.14 (0.15) | 0.12 | -0.16 to 0.44 | 1 | -0.07 (0.01) | -0.82 | -0.10 to -0.04 | **<0.001** |
| Acc 06:00-09:00 | -0.5 (0.09) | -0.43 | -0.68 to -0.33 | **<0.001** | -0.01 (0.01) | -0.06 | -0.02 to 0.01 | 1 |
| Acc 09:00-12:00 | 0.03 (0.1) | 0.02 | -0.16 to 0.22 | 1 | 0.01 (0.01) | 0.08 | -0.01 to 0.02 | 1 |
| Acc 12:00-15:00 | -0.08 (0.1) | -0.07 | -0.26 to 0.11 | 1 | 0.01 (0.01) | 0.13 | -0.00 to 0.03 | 1 |
| Acc 15:00-18:00 | -0.04 (0.08) | -0.03 | -0.19 to 0.12 | 1 | 0.02 (0.01) | 0.19 | 0.00 to 0.03 | 0.263 |
| Acc 18:00-21:00 | 0.18 (0.07) | 0.15 | 0.05 to 0.31 | 0.134 | 0.02 (0.01) | 0.22 | 0.01 to 0.03 | **0.015** |
| Acc 21:00-00:00 | -0.07 (0.08) | -0.06 | -0.22 to 0.08 | 1 | 0 (0.01) | -0.03 | -0.02 to 0.01 | 1 |
| Acc 00:00-03:00 | 0.26 (0.11) | 0.23 | 0.05 to 0.47 | 0.199 | -0.01 (0.01) | -0.13 | -0.03 to 0.01 | 1 |
| Acc 03:00-06:00 | -0.02 (0.09) | -0.01 | -0.19 to 0.16 | 1 | 0 (0.01) | -0.03 | -0.02 to 0.01 | 1 |
| Light 06:00-09:00 | -0.11 (0.08) | -0.09 | -0.26 to 0.05 | 1 | -0.02 (0.01) | -0.21 | -0.03 to -0.01 | 0.105 |
| Light 09:00-12:00 | -0.31 (0.08) | -0.26 | -0.45 to -0.16 | **0.001** | -0.01 (0.01) | -0.08 | -0.02 to 0.01 | 1 |
| Light 12:00-15:00 | 0.02 (0.08) | 0.01 | -0.14 to 0.17 | 1 | 0 (0.01) | 0 | -0.01 to 0.01 | 1 |
| Light 15:00-18:00 | 0.08 (0.07) | 0.07 | -0.06 to 0.23 | 1 | 0.01 (0.01) | 0.06 | -0.01 to 0.02 | 1 |
| Light 18:00-21:00 | -0.1 (0.08) | -0.09 | -0.25 to 0.05 | 1 | 0 (0.01) | -0.05 | -0.02 to 0.01 | 1 |
| Light 21:00-00:00 | -0.13 (0.08) | -0.11 | -0.28 to 0.02 | 1 | 0.01 (0.01) | 0.14 | -0.00 to 0.03 | 0.702 |
| Light 00:00-03:00 | 0.24 (0.1) | 0.21 | 0.04 to 0.44 | 0.248 | -0.01 (0.01) | -0.17 | -0.03 to 0.00 | 1 |
| Light 03:00-06:00 | 0.05 (0.08) | 0.04 | -0.11 to 0.21 | 1 | 0 (0.01) | -0.01 | -0.01 to 0.01 | 1 |
| Observations | 169 |  |  |  | 169 |  |  |  |
| R2 / R2 adjusted | 0.722 / 0.691 |  |  |  | 0.636 / 0.595 |  |  |  |

Abbreviations: B (SE) = unstandardized coefficient (standard error); $\beta$ =standardized coefficient; CI = 95% confidence interval. Predictors were weighted according to the type of day and standardized. Bold-formatted $p$ -values indicate statistical significance after Holm-Bonferroni correction.

**Figure 2.**
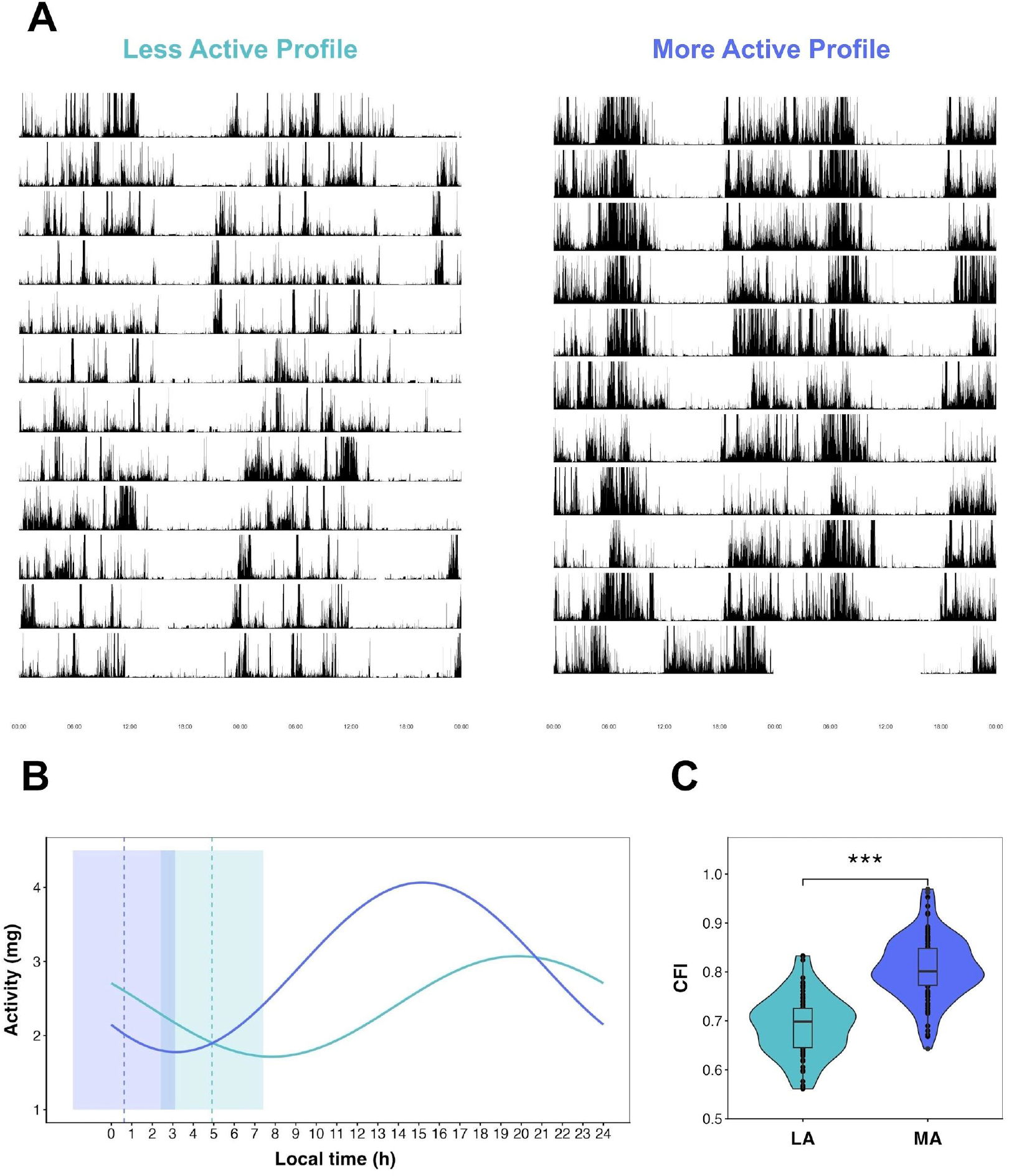
(a) Representative actograms from two participants belonging to different movement profiles. (b) Cosinor-fitted model (lines) and L5c (bars and vertical lines) for the same two participants. (c) MA showed higher CFI values compared with LA. The comparison was performed using the Wilcoxon rank-sum test. Statistical significance after Holm-Bonferroni correction is indicated as \**p* < 0.05, \*\**p* < 0.01, \*\*\**p* < 0.001.

**Figure 3.**
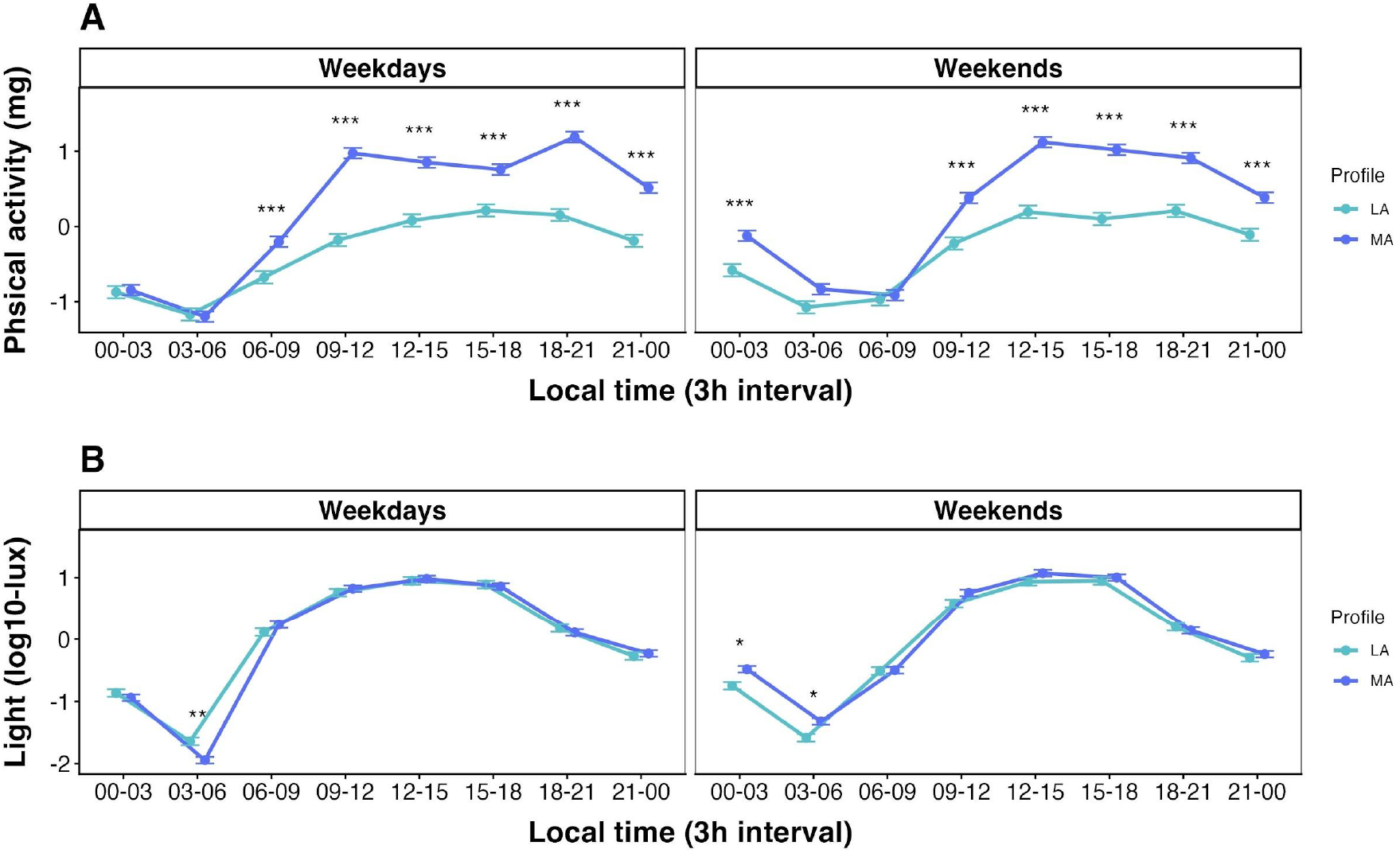
Scaled mean activity (a) and light exposure (b) across the day, shown in 3-hour intervals by profile and type of day. Both profiles showed different behavior regarding daily physical activity and light exposure. Statistical significance after Holm-Bonferroni correction is indicated as \**p* < 0.05, \*\**p* < 0.01, \*\*\**p* < 0.001.

### Effect size

We conducted an a priori power analysis to determine the minimum detectable effect sizes for planned analyses. For comparisons between two independent groups, we used the pwr.t.test() function from the pwr package in R (Champely, 2006), assuming equal group sizes and a two-tailed test (α=0.05, power=0.9). Under these conditions, a minimum of 84 participants per group would allow detection of an effect size of d≈0.5, considered moderate according to Cohen’s benchmarks. The actual profiles resulting from clustering included 96 participants in one profile and 73 in the other. A post hoc power analysis using pwr.t2n.test() indicated that the achieved power for detecting d ≈ 0.5 was also 0.9. For the multiple regression models (17 predictors, *n* = 169, α = 0.05, power = 0.9), an a priori power analysis using pwr.f2.test() indicated an expected f^2^ ≈ 0.16, corresponding to R^2^ ≈ 0.14.

## RESULTS

### Movement profiles

The k-means algorithm identified two distinct clusters derived from 12 accelerometer-based movement features: the Less Active profile (LA, *n* = 73) and the More Active profile (MA, *n* = 96). Although the average silhouette widths were modest (LA: 0.34, MA:0.33), the two clusters did not overlap and fulfilled the goal of capturing dominant behavioral dimensions. (Figure 1). Interestingly, these two profiles that emerged from an unsupervised machine learning method showed no differences in sex, age, BMI or chronotype between them (73% female in both profiles, LA: 22.3 ± 3.1 years, BMI=24.3±3.6 Kg/m^2^, MSFsc = 05:38 ± 01:51; MA: 23.5 ± 3.4 years, BMI=23.8±3.5, MSFsc = 05:15 ± 01:45, *p*>0.05). But, consistent with the parameters that fed the model, 11 out of the 12 movement features exhibited significant differences between MA and LA after Holm-Bonferroni correction (Table 2).

Actigraphy was recorded for a period of 16.5 ± 4.2 (range 8-24) days across all participants, with 15.2 ± 4.1 (range 8-23) days in LA and 17.4 ± 4.0 (range 8-24) days in MA. The proportion of weekends over weekdays was 0.51 ± 0.12 overall, 0.52 ± 0.13 in LA, and 0.50 ± 0.12 in MA. Day length was 12.06 ± 0.65 (10.96-13.27) hours overall, 12.32 ± 0.47 (10.96-13.25) in LA, and 11.86 ± 0.70 (10.96-13.27) in MA. Cohorts were distributed heterogeneously across the two profiles (see Supplementary Table S1). The 2019 dancers cohort was predominantly MA (93%), the 2021 dancers cohort was 71% MA, the 2016 university cohort was largely LA (69%), and the 2014 university cohort, the largest, was 44% LA.

### Sleep and circadian rhythm parameters between profiles

L5c, M10c and acrophase were significantly associated with MSFsc (β = 0.69, 0.45, and 0.6 respectively; all *p*<0.001), independent of sex and age. L5c has been shown to correlate with physiological measures of circadian phase (dim-light melatonin onset, DLMO) in previous studies with the same cohorts, supporting its use as a proxy for circadian phase (Castillo et al., 2023; Coirolo et al., 2022). We observed no significant differences in parameters associated with sleep quality between MA and LA (Table 3). MA exhibited an earlier L5c with a small effect size; however, neither M10c nor the acrophase differed between profiles In contrast, all rhythm-related features showed significant differences: MA showed higher amplitude and mesor (estimated by the cosinor model), as well as more robust (higher RA), less fragmented (lower IV), and more regular (higher IS) rest-activity rhythms (Table 3). These characteristics are clearly visible in the representative actograms of MA and LA presented in Figure 2a. Furthermore, the overall integrity of the rhythms, summarized by the CFI, differed significantly between profiles, with MA showing higher values (0.81 ± 0.06 versus 0.69 ± 0.06, *p*<0.001, effect size = 0.697) (Figure 2b).

### Physical activity and light exposure by movement profile and type of day

To explore how movement profiles relate to behavior throughout the day, we examined physical activity and light exposure across 3-hour intervals on weekdays and weekends. Mixed models revealed significant between-profile differences in the marginal means of physical activity across intervals for most times of day, with exceptions at 00:00_-_03:00 and 03:00_-_06:00 on weekdays, and 03:00_-_06:00 and 06:00_-_09:00 on weekends (Figure 3A). In contrast, for light exposure, significant differences occurred only at 00:00_-_03:00 on weekdays, and 00:00_-_03:00 and 03:00_-_06:00 on weekends (Figure 3b).

### Associations of circadian phase and overall circadian rhythms integrity with daily physical activity and light exposure across time-of-day intervals

L5c was negatively associated with early morning physical activity (06:00_-_09:00) and late morning light exposure (09:00_-_12:00) after Holm-Bonferroni correction (Table 4). No other activity or light exposure intervals, nor the movement profile, were significantly associated with L5c. Overall integrity of the circadian activity–rest rhythm, estimated by CFI, was positively associated with physical activity during the early evening (18:00–21:00) and negatively associated with LA (Table 4). No other activity or light exposure intervals remained significantly associated after correcting for multiple comparisons. Sex, age, and day length were evaluated as covariates but were not retained in the final models because their inclusion did not improve model fit (AIC) and none showed statistically significant effects.

CFI showed a small negative association with L5c (B=-0.01, SE=0.01, p=0.035); however, this effect became non-significant after adjusting for sex and age and including the movement profile in the model (B=0, SE=0.01, p=1), with belonging to LA remaining strongly associated with lower CFI (B=-0.12, SE=0.01, p<0.001) after Holm-Bonferroni correction. Finally, CFI exhibited a significant association with the Sleep Regularity Index (B = 3.53, SE=1.17, *p* = 0.012) and sleep duration (B=0.22, SE=0.08, *p* = 0.019), but not with WASO or SE (p>0.05).

## DISCUSSION

A central challenge in circadian health research is the identification of accessible and reliable markers that capture the integrity of the human circadian rhythms in real-world settings. To our knowledge, this is the first work using unsupervised machine-learning to detect movement profiles and assess their association with circadian rhythms integrity in healthy young adults in real-life conditions. The clusters seem to capture meaningful biological variation, as shown by consistent differences in independent circadian rhythm metrics. Consistent with our hypothesis, MA participants showed healthier activity-rest circadian rhythm (i.e., higher CFI) than LA participants. Our findings highlight the relevance of movement profiles in two ways: as predictors of circadian rhythm health, and as a way to re-entrain misaligned rhythms.

In the broader effort to integrate circadian concepts into medicine and clinical practice, simple, cost-effective, and scalable metrics are essential. Our study contributes evidence of non-invasive markers of circadian health, alongside novel methodological approaches designed to streamline assessment and reduce cost and time, in line with recent developments (Gubin et al., 2025; Huang et al., 2021; Lim et al., 2024, 2025). Perhaps the most notable finding of this study is the association between movement profiles and overall circadian rhythms integrity, as assessed by the CFI. This index was able to capture relevant information from real-world data by combining circadian key parameters into a single and easily interpretable value (Ortiz-Tudela et al., 2010). Although the CFI is not yet clinically validated for diagnostic purposes, it could be considered analogous to general health indexes in medicine, such as cholesterol levels in cardiometabolic health.

Large epidemiological studies have linked lower RA to adverse health outcomes, including increased all-cause mortality and higher incidence of cardiovascular disease, cancer, and infectious, respiratory, and digestive disorders (Feng et al., 2023). Most evidence comes from predominantly White adult cohorts, highlighting the need for research in more diverse populations. In our study, a more active behavioral profile was associated with greater RA in healthy, young higher-education students from an admixed population. While causal relationships cannot be inferred, these findings underscore the role of physical activity in supporting circadian robustness and overall health.

Disruptions of circadian rhythmicity contribute to disease development and premature aging. Fragmented, irregular, and weak rhythms have been associated with several health issues (Van Someren and Riemersma-Van Der Lek, 2007). A bidirectional link between circadian perturbations and oxidative stress has been proposed as a potential underlying mechanism (Drăgoi et al., 2024). In contrast, structured physical activity appears to modulate redox homeostasis, depending on intensity, frequency, and duration (McClean and Davison, 2022). In our study, MA displayed a richer and more complex physical activity pattern, as evidenced by higher overall acceleration, longer and more frequent LPA and MVPA bouts, reduced inactivity bout duration, and a markedly steeper intensity gradient, indicating engagement across a broader spectrum of intensities throughout the day. This enhanced activity pattern was associated with more stable, less fragmented, and more robust circadian rhythms. In addition, MA exhibited higher amplitude and mesor, parameters that are also positively associated with health and well-being.

Although DLMO remains the gold standard for estimating circadian phase, its routine clinical use is limited by cost and logistical constraints. This study adds evidence to the use of L5c as an alternative accelerometer-base phase estimator in healthy young populations (Bonmati-Carrion et al., 2014; Castillo et al., 2023; Coirolo et al., 2022). However, its reliability may be reduced in older or clinically challenged populations. Rather than a limitation, this provides an interesting perspective on its potential as a complementary marker of circadian health.

The influence of zeitgebers varies throughout the day, with specific sensitive windows in which their effect on circadian phase occur (Burgess et al., 2002). Bright light exposure induces phase delays at evening and night, and phase advances at early-morning (Khalsa et al., 2003), while exercise induces phase delays at evening and phase advances at both early-morning and early-afternoon (Youngstedt et al., 2019). Although a small difference in L5c was noted among profiles, disaggregating this by time-of-day revealed associations with light exposure and exercise timing, irrespective of profile. Both morning light exposure and morning exercise were associated with a tendency to earlier L5c values. In contrast, CFI did not correlate with light exposure at any time of day, suggesting at least a partial dissociation between circadian phase and rhythm integrity.

The entrainment of skeletal muscle clocks through scheduled physical activity may enhance metabolic efficiency (Gabriel and Zierath, 2019). Afternoon and evening exercise appears to confer advantages for individuals with metabolic challenges (Shen et al., 2023), including lower post-exercise glucose levels in type 1 diabetes (Toghi-Eshghi and Yardley, 2019) and greater improvements in glycemic control following high-intensity interval training in men with type 2 diabetes (Savikj et al., 2019). These benefits may arise from increased core and skeletal muscle temperatures, which enhance metabolic reactions, improve contractile efficiency, and boost neuromuscular activation (Gabriel and Zierath, 2019). Although the association between CFI and early evening physical activity in our study was modest, it aligns with evidence that afternoon and evening exercise may support circadian rhythm regulation.

Regarding sleep, although circadian and homeostatic processes are related, it is important to distinguish between them, as they have different implications for health (Borbély, 2022). In our study, MA exhibited better outcomes in several circadian metrics, but no differences were observed between profiles concerning sleep quality indicators. This is a noteworthy result, as it suggests that sleep homeostasis and circadian rhythmicity can be disentangled when examined through the lens of movement behavior, consistent with recent CFI-based studies distinguishing circadian function from sleep duration (Samson and McKinnon, 2025). Nevertheless, higher CFI values were associated with better sleep quality, regardless of the movement profile.

Lower CFI values, reflecting weakened circadian rhythms, have been observed in individuals with a variety of health issues, including Parkinson’s disease and sleep-disordered breathing, compared with controls (Madrid-Navarro et al., 2018; Martinez-Nicolas et al., 2021). This circadian dysfunction, resulting from either aging or illness, requires strong and regularly timed zeitgebers to achieve more robust rhythms (Van Someren and Riemersma-Van Der Lek, 2007). While exposure to effective light-dark contrast is difficult in urban environments, timed physical activity offers a promising alternative as a practical strategy to support circadian alignment and overall health. For instance, reducing sedentary behavior and increasing activity has been recently associated with stronger rest–activity rhythms in cancer survivors (Frenken et al., 2025). A conceptually similar approach integrating multiple circadian rhythm dimensions has been applied in two large cohorts of predominantly White older adults (Vidil et al., 2025). Notably, despite differences in demographic profiles, the correlation structure among circadian rhythm dimensions appeared largely consistent (Figure SM1). By including the CFI, our approach highlights additional aspects of circadian health integrity.

Our study addresses both the strengths and limitations of ecological models. We did not account for the menstrual cycle, nor were able to identify the specific types of physical activity participants engaged in or control for other factors such as food habits or social pressures. In addition, actigraphy has inherent limitations, including issues with body placement, which may affect both the accuracy of movement detection by the accelerometer and light exposure measurement, due to the distance of the light sensor from the eye, and potential obstruction by clothing (Gubin et al., 2025). Although merging databases increased sample size for actigraphy, this approach also resulted in some data being missing or incomplete, and potential confounding effects. Actimetry assessments were conducted around peri-equinox periods; however, day length differed between cohorts, which also varied in their habitual physical activity patterns. While cohort heterogeneity—from dancers to more sedentary participants—may be considered a limitation, it strengthens the analysis by capturing diverse circadian behaviors. Despite these limitations, our results provide valuable insights into the relationship between movement behavior and overall circadian rhythms integrity. Future research should include longitudinal studies, explore different age groups and health conditions, compare real-world, lab and clinical settings, distinguish types of physical activity, and incorporate additional factors such as diet and social influences.

## CONCLUSION

Three aspects stand out as the main contributions of this study. First, it reinforces the value of actimetry as a standalone objective measure by exploiting its multidimensional nature, which can provide clinically and scientifically meaningful insights. Second, by summarizing rhythmic strength, regularity, and fragmentation, the use of CFI offers a practical way to incorporate circadian information into clinical research and potentially into future screening or monitoring strategies. Third, our findings suggest that individuals with a more active physical activity profile exhibit stronger overall circadian rhythms integrity, which has been linked to better health outcomes. Overall, our work offers a replicable integrative framework based on movement behavior as a core dimension of circadian health, spanning educational, cultural, and clinical contexts.

## ACKNOWLEDGMENTS

We specially thank the participants for kindly taking part in this research. We also thank Julieta Castillo Stratta, Paloma Mombrú and Nicolás Parrillo for their collaboration in new data collection, and Natalia Coirolo for her contribution in previous work. We also thank the staff of Facultad de Ciencias for their assistance, and the members of the Grupo Cronobiología for their feedback at different stages of the process.

## AUTHORS CONTRIBUTION

Conceptualization and writing–original draft: MM, AS and BT; formal analysis, MM; funding acquisition and supervision AS and BT. All authors have read and agreed to the final version of the manuscript to be published.

## AVAILABILITY OF DATA AND MATERIALS

Data will be available from the corresponding author upon reasonable request.

## ETHICAL CONSIDERATIONS AND CONSENT TO PARTICIPATE

This study was conducted according to the guidelines laid down in the Declaration of Helsinki and all procedures involving human subjects were approved by the Ethics Committee of the School of Psychology, Universidad de la República (Nº 191175-000124-23). All participants provided written informed consent prior to participation.

