## Supplementary material for "Actigraphy-Based Movement Profiles and Their Association With Circadian Rhythms Integrity in Real-World Settings"

**Table S1** Baseline characteristics of the study cohorts, including seasonal context, demographics, profile proportions, physical activity measures used for clustering, and circadian and sleep quality parameters.

| | **New data** | **Marchesano et al., 2025** | **Coirolo et al., 2022** | **Silva et al., 2019** |
|---|---|---|---|---|
| | N = 95 | N = 31 | N = 30 | N = 13 |
| | University students | Dancers | Dancers | University students |
| **Age** | 23.4 (3.6) | 22.5 (3.4) | 22.4 (2.8) | 22.8 (1.5) |
| **Sex (Female)** | 65% (62) | 87% (27) | 83% (24) | 69% (9) |
| **BMI** | 24.7 (3.6) | 23.6 (3.6) | 23.1 (2.2) | 21.8 (3.4) |
| **Chronotype (MSFsc)** | 5.23 (1.93) | 5.22 (1.38) | 6.13 (1.85) | 5.95 (0.84) |
| **Actimetry assessment (month–year)** | Sep–Oct 2024 | Sep 2021 | Sep 2019 | Mar 2016 |
| **Day length (h)** | 12.15 (0.46) | 11.48 (0.01) | 10.67 (0.00) | 12.57 (0.00) |
| **Profile (MA)** | 44% (42) | 71% (22) | 93% (28) | 31% (4) |
| **Average daily acceleration (mg)** | 44 (12) | 51 (15) | 56 (11) | 42 (11) |
| **Total duration in IN (min/day)** | 694 (76) | 657 (97) | 600 (64) | 685 (92) |
| **Total duration in LPA (min/day)** | 171 (39) | 180 (44) | 196 (33) | 167 (43) |
| **Total duration in MVPA (min/day)** | 120 (47) | 155 (66) | 177 (49) | 111 (56) |
| **Mean duration of IN bouts (min)** | 14.8 (4.5) | 13.1 (6.4) | 9.8 (2.0) | 13.5 (3.8) |
| **Mean duration of LPA bouts (min)** | 2.56 (0.29) | 2.46 (0.25) | 2.44 (0.21) | 2.45 (0.35) |
| **Mean duration of MVPA bouts (min)** | 3.61 (0.92) | 3.81 (0.79) | 4.12 (0.76) | 3.19 (0.95) |
| **Daily bouts in IN (n)** | 52 (11) | 59 (14) | 63 (10) | 55 (11) |
| **Daily bouts in LPA (n)** | 61 (14) | 68 (17) | 75 (12) | 62 (14) |
| **Daily bouts in MVPA (n)** | 33 (10) | 39 (12) | 44 (8) | 33 (10) |
| **Intensity gradient** | -1.89 (0.14) | -1.86 (0.15) | -1.80 (0.09) | -1.91 (0.16) |
| **Intensity intercept** | 11.86 (0.45) | 11.76 (0.48) | 11.65 (0.36) | 11.89 (0.56) |
| **Sleep Duration (h)** | 6.52 (0.71) | 6.44 (0.88) | 6.59 (0.63) | 6.63 (1.04) |
| **Sleep Regularity Index** | 55 (11) | 50 (12) | 50 (10) | 58 (10) |
| **WASO (h)** | 1.07 (0.38) | 1.06 (0.40) | 1.09 (0.38) | 1.27 (0.29) |
| **Sleep Efficiency (%)** | 86.2 (4.8) | 86.2 (4.3) | 86.4 (4.1) | 84.5 (3.5) |
| **L5c (local time)** | 26.14 (1.26) | 26.26 (0.81) | 26.27 (1.23) | 26.63 (0.99) |
| **M10c (local time)** | 11.23 (1.45) | 10.87 (1.23) | 11.50 (2.16) | 11.62 (1.86) |
| **Acrophase (cosinor, rad)** | 4.34 (0.36) | 4.25 (0.24) | 4.34 (0.43) | 4.48 (0.36) |
| **Amplitude (cosinor)** | 0.88 (0.17) | 0.89 (0.21) | 0.87 (0.18) | 0.87 (0.14) |
| **Mesor (cosinor)** | 2.63 (0.17) | 2.66 (0.21) | 2.74 (0.16) | 2.62 (0.18) |
| **Relative Amplitude (RA)** | 0.84 (0.04) | 0.86 (0.05) | 0.88 (0.02) | 0.82 (0.05) |
| **Interdaily Stability (IS)** | 0.33 (0.10) | 0.33 (0.12) | 0.34 (0.09) | 0.36 (0.07) |
| **Intradaily Variability (IV)** | 0.96 (0.17) | 0.81 (0.13) | 0.81 (0.13) | 1.08 (0.14) |
| **CFI** | 0.74 (0.09) | 0.79 (0.08) | 0.80 (0.06) | 0.70 (0.07) |

Mean (SD); % (n)

**Figure S1.** Correlation matrix reflecting movement features, sleep quality and circadian parameters. Only correlations that remained statistically significant after Holm correction for multiple comparisons ($\alpha$ = 0.05) are displayed; non-significant associations are left blank. Correlation coefficients are reported in black within each cell. The diagonal represents self-correlations.

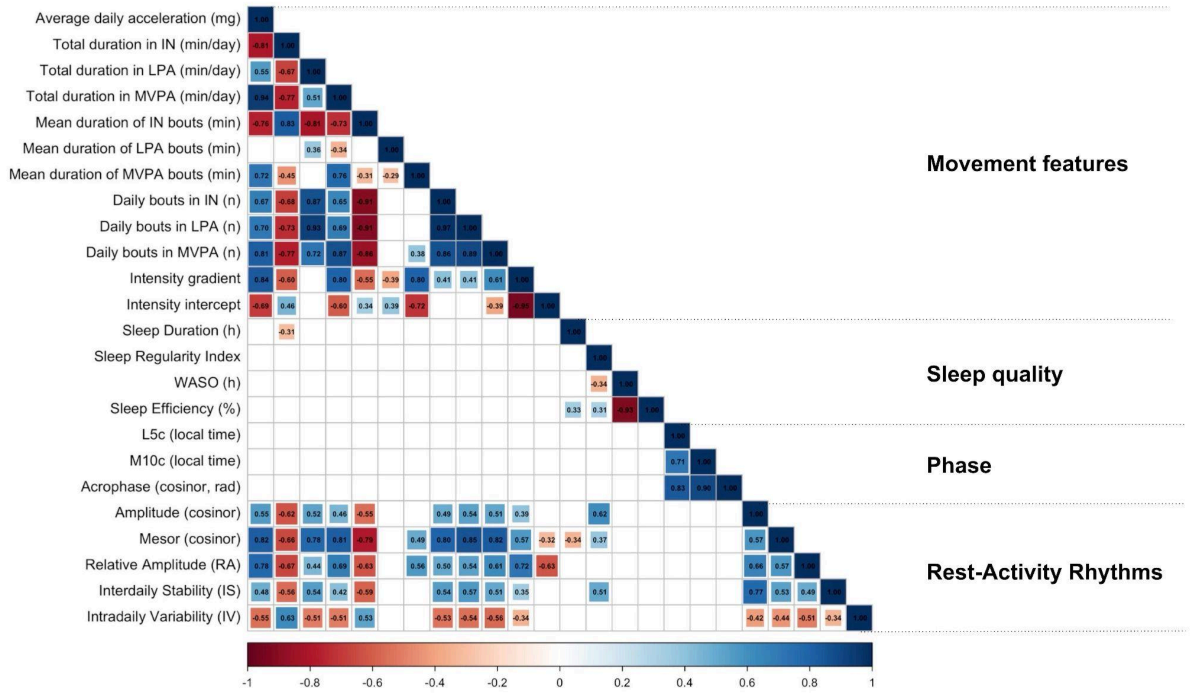
